# Semi-Analytic Nonparametric Bayesian Inference for Spike-Spike Neuronal Connectivity

**DOI:** 10.1101/340489

**Authors:** Luca Ambrogioni, Patrick W. J. Ebel, Max Hinne, Umut Güçlü, Marcel A. J. van Gerven, Eric Maris

## Abstract

Estimating causal connectivity between spiking neurons from measured spike sequences is one of the main challenges of systems neuroscience. In this paper we introduce two nonparametric Bayesian methods for spike-membrane and spike-spike causal connectivity based on Gaussian process regression. For spike-spike connectivity, we derive a new semi-analytic variational approximation of the response functions of a non-linear dynamical model of interconnected neurons. This semi-analytic method exploits the tractability of GP regression when the membrane potential is observed. The resulting posterior is then marginalized analytically in order to obtain the posterior of the response functions given the spike sequences alone. We validate our methods on both simulated data and real neuronal recordings.

## 1 Introduction

Action potentials (spikes) are the fundamental units of neuronal communication [1]. Spikes originate from the axon hillock and propagate through the axon towards the synaptic terminal, where the release of neurotransmitters affects the membrane potential of the downstream neurons. While there is a great deal of computation in the dynamics of a single neuron [2], most of the computational capabilities of biological neuronal networks depend on their pattern of interconnections [3]. Mapping causal interrelations between spiking neurons is therefore a major goal in system neuroscience. However, inferring causal connectivity from spike sequences is a challenging data analysis problem as networks of spiking neurons are highly non-linear dynamical systems [4].

In this paper we introduce two related methods for the estimation of the causal response function between spiking neurons based on Gaussian process (GP) regression. Both methods rely on an important neurophysiological fact concerning neuronal communication: the membrane potential responds approximately linearly to weak synaptic inputs while spike initiation is a highly non-linear function of the membrane potential [5, 6]. The first of our new methods is applicable when both spike sequences and membrane potentials are observed variables. In this case, the posterior distribution of the resulting connectivity model can be obtained analytically and is related to the GP-CaKe method for field-field causal connectivity [7]. The main methodological contribution of the paper is in our second method, which is applicable when only the spike sequences are measured. In this situation, the Bayesian model cannot be solved in closed form since the spike initiation model is non-Gaussian. Furthermore, approximate inference is complicated by the intractability of the inhomogeneous Poisson likelihood [8, 9]. To resolve these difficulties, we derive a new semi-analytic variational approximation that combines the analytic solution of the response function given the membrane potential with a likelihood-free stochastic estimation of the membrane potential.

## 2 Spike-membrane causal connectivity analysis

In this section we introduce a nonparametric Bayesian method for estimating the causal response function when both spike sequences and membrane potentials are observed variables. Besides its intrinsic relevance in several experimental settings, this method is also an important analytically tractable component of our method for spike-spike connectivity. We begin by introducing a linear dynamical model of the membrane potential that captures the linear response of the membrane potentials to weak synaptic inputs.

### 2.1 A linear dynamical model of the membrane potential

Consider a network of *N* interconnected neurons. In the following, we will denote the membrane potential of the *j*-th neuron as *m*_*j*_ (*t*) and its spike sequence as the sum of delta functions *s*_*k*_ (*t*) = Σ_*k*_ *δ*(*t* − *t*_*j*,*k*_) where *t*_*j*,*k*_ is the timestamp of the *k*-th spike of the *j*-th neuron. The linear response of a neuronal membrane to a synaptic input can be described using a differential equation [7]:

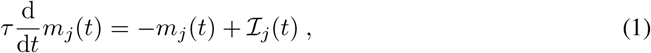

where the time constant *τ* determines the time that the membrane needs to return to baseline after a perturbation. The synaptic input from the other *N* − 1 neurons in the network is given by the following function:

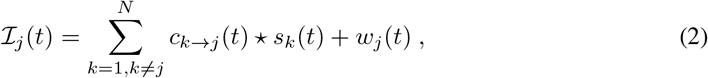

where the operator ★ denotes convolution. The additional stochastic term *w*_*j*_(*t*) is Gaussian white noise with variance *σ*^2^ and accounts for unmeasured perturbations. The causality of the neuronal network is guaranteed as the causal response function *c*_*k*→*j*_(*t*) vanishes for negative values of *t*.

### 2.2 Analytic GP regression for spike-membrane causal connectivity

We use the dynamical model specified by Eq. 1 and Eq. 2 as an implicit likelihood of a nonparametric Bayesian model. The model is defined by assigning a GP prior over the space of response functions *c*_*k*→*j*_. The posterior distribution of *c*_*k*→*j*_ is a GP and can be obtained in closed-form because both the derivative and the convolution in Eq. 1 and Eq. 2 are linear operators. In the frequency domain, Eq. 1 and Eq. 2 can be jointly written as

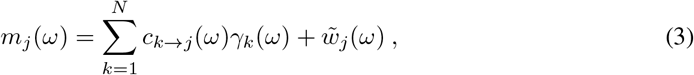

where

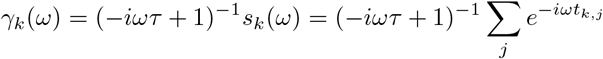

and

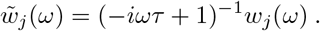

Eq. 1 defines a nonparametric regression problem where *m*_*j*_ (*ω*) is the observed data, *γ*_*k*_ (*ω*) are known mixing functions and *c*_*k*→*j*_(*ω*) are the unknowns. Problems of this form have an analytic solution when the prior distributions over *c*_*k*→*j*_(*ω*) are GPs [10]. To assure the causality of the response functions we adopt the causal covariance function that was introduced in [7]. In the frequency domain this covariance function can be expressed as

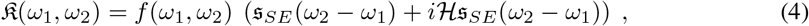

where *f*(*ω*_1_ *ω*_2_) is a function that induces smoothness by discounting the high frequency components, 𝔰_*SE*_(*ω*) is the spectral density of a squared exponential covariance function and 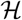 denotes the Hilbert transform which enforces causality. The resulting GP prior induces causality, smoothness and temporal localization of the response function. See Appendix A for more details on the construction of this covariance function.

Consider a set of *M* time points {*t*_1_, …,*t*_M_} and a vector of measured membrane potentials *m*_*u*_ = *m*(*t*_*u*_). The posterior expected value of *c*_*k*→*j*_ is given by

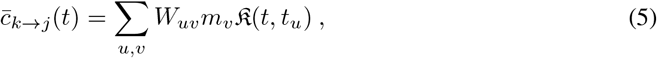

where the GP weights *W*_*uν*_ depend on the covariance function and can be obtained using standard GP regression techniques in the frequency domain. The matrix formula for the weights is given in Appendix B. The time domain covariance function in Eq. 5 is the inverse Fourier transform of Eq. 4 with respect to both of its arguments.

## 3 Spike-spike causal connectivity analysis

We can now use the results of the previous section in order to derive a semi-analytic solution to the more challenging problem of spike-spike connectivity.

### 3.1 A non-linear model of spike initiation

In biological neurons, spike initiation depends on the non-linear dynamics of the membrane potential and of several ionic channels [4]. We approximate these dynamics using a stochastic model. Specifically, the firing rate *f*(*t*) is obtained by passing the rescaled membrane potential through a compressive non-linearity:

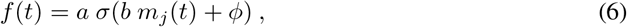

where *a* is the maximum firing rate and *σ*(·) is the logistic sigmoid with *b* and *ϕ* its gain and threshold parameters respectively. The resulting spike sequence follows a nonhomogeneous Poisson process with density function *f*(*t*) [8]. This model is admittedly a simplification. For example, it does not take into account the refractory period [11]. However, the variational Bayesian model that we will introduce in the next section can be used with any other spike initiation model without substantial modifications.

### 3.2 Semi-analytic variational GP regression for spike-spike causal connectivity

To simplify the notation we will explain the analysis for the case of two neurons. All results generalize straightforwardly to arbitrary network structures. Given a set of *M* sample time points {*t*_1_,…,*t*_*M*_}, we organize the sampled time-series in the arrays *s*_*j*_ = (*s*_*j*_ (*t*_1_,‥,*s*_*j*_(*t*_*M*_)), ***m***_*j*_ = (*m*_*j*_(*t*_1_),‥, *m*_*j*_(*t*_*M*_)) and *c*_2→1_ = (*c*_2→1_(−*t*_*M*_/2), ‥, *c*_2→1_(*t*_*M*_/2)). The graphical model is shown in Fig. 1. This model is summarized by the following factorized joint distribution:

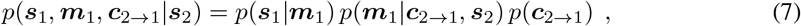

where we conditioned on the spike sequence *s*_2_. Our aim is to obtain *p*(*c*_2→1_|*s*_1_,*s*_2_),i.e., the posterior distribution of the causal response function given the two spike sequences. Most existing variational methods do not directly leverage the analytic solution of *p*(*c*_2→1_|*m*_1_,*s*_2_) and require the evaluation of the intractable likelihood *p*(*s*_1_|*m*_1_)[12–14]. Therefore we developed a new semianalytic variational approximation that fully exploits the analytic tractability of the latent GP analysis. We begin by defining the following structured joint variational distribution:

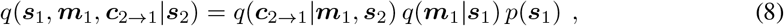

where *p*(*s*_1_) is the real marginal distribution of *s*_1_. In this variational factorization we assumed that the distribution of the membrane potential *m*_1_ solely depends on the spike sequence *s*_1_. We can find the distributions *q*(*c*_2→1_|*m*_1_, *s*_2_) and *q*(*m*_1_|*s*_1_) by minimizing the following functional:

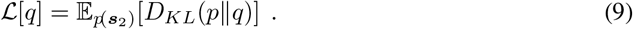

Note that this functional is a proper (joint-contrastive) variational loss since it is always non-negative and vanishes if and only if *p*(*s*_1_,*m*_1_,*c*_2→1_,*s*_2_ = *q*(*s*_1_,*m*_1_,*c*_2→1_,*s*_2_).We can rearrange the loss as:

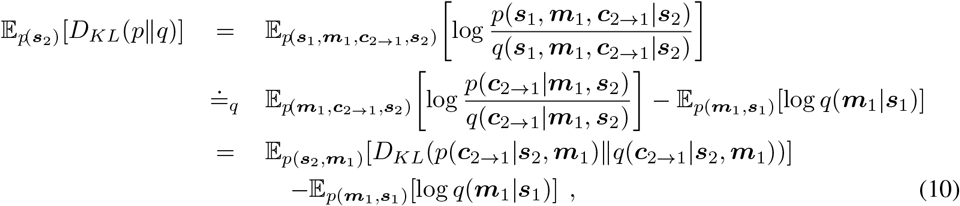

where ≐_*q*_ denotes that the expressions are equal up to terms that are constant in *q*. The first term of this expression is an expectation of a KL divergence and therefore vanishes when *q*(*c*_2→1_ | *s*_2_,*m*_1_) is equal to the real posterior *p*(*c*_2→1_|*s*_2_,*m*_1_), which can be expressed analytically (see Eq.5).We can parameterize the remaining term as a mixture of Gaussian distributions:

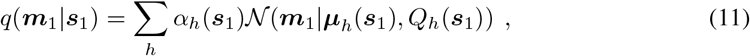

where the scalar-valued functions α_h_(*s*_1_), the vector-valued functions ***μ***_h_(*s*_1_) and the matrix-valued functions *Q*_*h*_(*s*_1_) are determined by expressive regression models such as deep convolutional networks [15]. The parameters of these networks can be trained by minimizing the remaining term of the variational loss

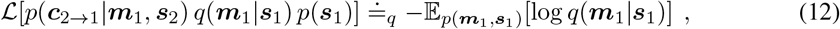

whose gradient can be easily sampled without bias by sampling from the model marginal *p*(*m*_1_,*s*_1_). Optimizing Eq.12 requires to train the regression models *α*_*h*_(*s*_1_), ***μ***_*h*_(*s*_1_) and *Q*_*h*_(*s*_1_) separately every time we want to analyze a new network structure since the distribution *p*(*m*_1_; *s*_1_) includes the (marginalized) effects of all neurons. In order to increase the efficiency of the method we approximate *p*(*m*_1_; *s*_1_) with the joint distribution of a single uncoupled neuron. This is a weak coupling approximation since we are assuming the (cumulative) coupling strength between neurons to be small compared to the stochastic input. We analyze the consequences of this approximation in our experiments below. We can now obtain the variational posterior *q*(*c*_2→1_|*s*_1_, *s*_2_) by marginalizing the variational distribution analytically:

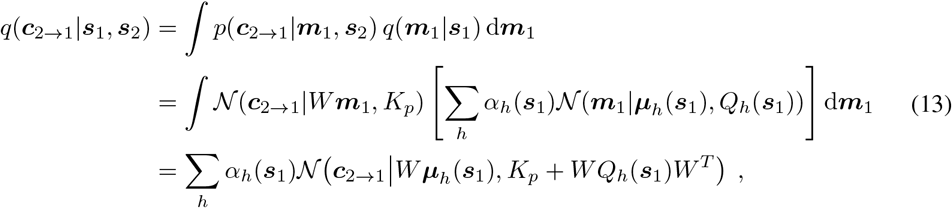

where *W* are the GP weights (see Eq. 5) and *K*_*p*_ is the covariance matrix of the posterior *p*(*c*_2→1_|*m*_1_, *s*_2_). We refer to our method of spike-spike connectivity estimation as SGP CaKe (Spike Gp Causal Kernels).

**Figure 1:**
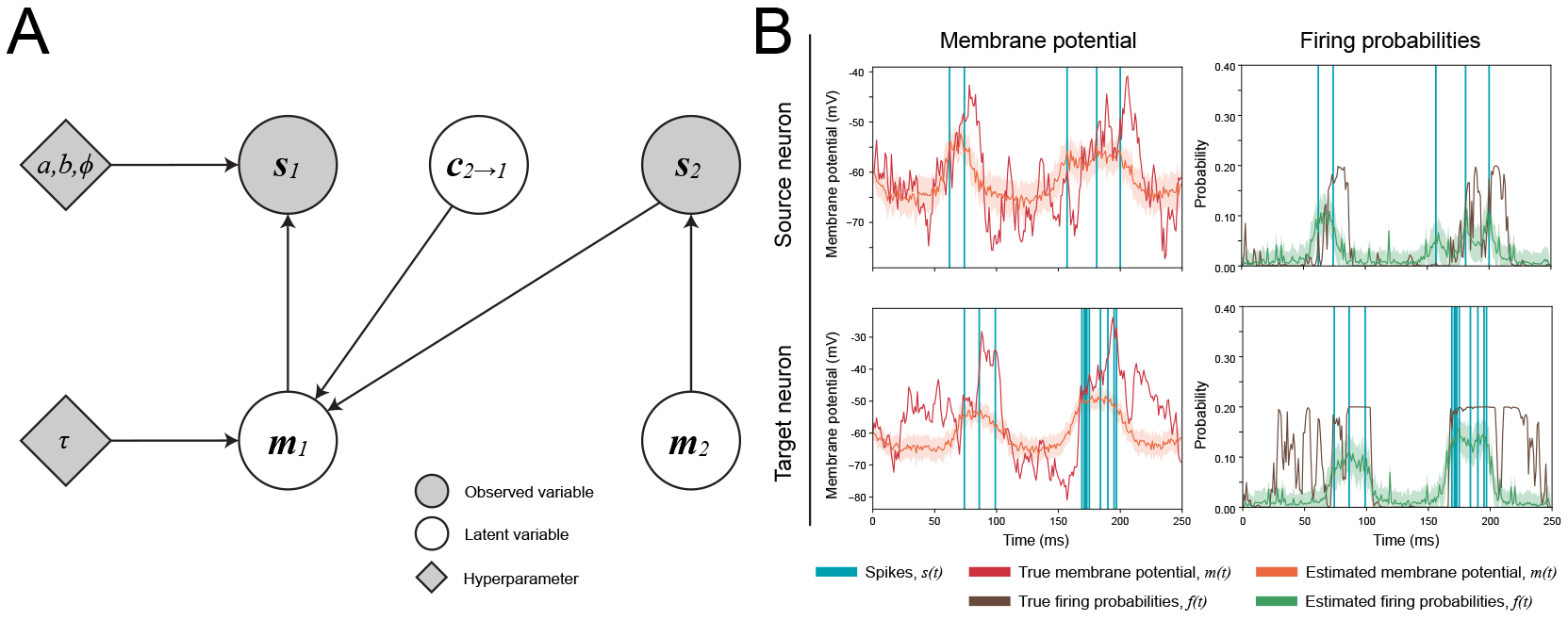
**A.** The generative model as explained in the text. For simplicity of the notation, the relevant variables are only shown for two neurons. Note that the membrane potentials *m* can be either observed or latent. In the latter case we use the variational approach. **B.** A draw from the generative model for two neurons connected through a single unidirectional excitatory connection as well as the variational recovery of the membrane and action potentials. The shaded regions indicate one standard deviation.

## 4 Related work

Several techniques have been used to identify spike-spike connectivity. Simple nonparametric methods such as histograms have a long history and are still widely applied [16]. Parametric methods based on the generalized linear model (GLM) often offer a better signal-to-noise ratio [17]. The models introduced in this paper are strictly related to GP classification and can therefore be considered as the nonparametric generalization of GLM based methods [10]. Other modern approaches are based on dynamic Bayesian networks [18] and Cox processes [19]. We will now devote special attention to methods based on Hawkes processes, given their theoretical similarity to our approach.

### 4.1 Spike-spike connectivity with Hawkes Processes

The multivariate stochastic process defined in this paper has some similarity with a Hawkes process [20]. While most of the existing literature based on Hawkes processes assumes a simple parametrization for the response functions, several new studies introduced the use of nonparametric methods [21–23]. Hawkes processes have been successfully used in neuroscience settings in order to infer spike-spike causal connectivity [24, 25]. In a Hawkes process the spike density of the *j*-th unit is a linear functional of the spike sequences of the other units:

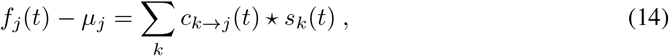

where *μ*_*j*_ is the baseline spike density. Note that Eq. 14 is strikingly similar to our Eq. 1. The difference is that in a Hawkes Process the spike density is a linear functional of the input spike sequences while in our model the linear response is defined at the level of the latent membrane potential. From a biophysical point of view, linearity of the spike density response is not a realistic assumption2019;threshold’ events [4]. Another obvious problem of Eq. 14 is that the spike density could become negative in the presence of inhibitory responses. Conversely, in our model a highly negative membrane potential simply corresponds to a very low but positive spike density. The similarity between Eq. 1 and Eq. 14 implies that both the analytic and the semi-analytic methods introduced in this paper can be applied to Hawkes processes as well. The analytic method cannot be applied on real data since the spike density is not directly measurable. Nevertheless we will use it as an idealized baseline comparison in our simulation studies where we know the ground truth.

## 5 Simulated effective connectivity

Here we validate the reconstructions by SGP CaKe. The details of the deep neural networks used for the estimation of the membrane are given in appendix E. The performance of spike-spike connectivity methods and non-linear regression in general is strongly affected by the form of regularization used. In order to have a balanced comparison we compare the performance of our method with its equivalent Hawkes process model where the prior covariance function and the approximative inference methods are exactly the same. We also include a comparison with a simpler nonparametric method based on spike-spike histograms [16].

First we define five different network structures, as shown in Fig. 4A. For each of these structures, which may contain both excitatory and inhibitory interactions, we generate 200 trials of observable membrane potentials and spikes according to the generative model of Section 3.2. The true connection strength w is varied to investigate its effect on the recovery of the causal response function. More details of the simulation procedure can be found in Appendix C. The first two networks simply demonstrate the recovery of either excitatory, inhibitory or absent coupling. An example of a single trial of simulated data is shown in Fig. 1B. The leftmost subfigures show the recovery of the membrane potential using the variational procedure. Note that as expected the spike density of neuron 2 is temporally concentrated near the spikes of the input neuron. Importantly, the transformation from membrane potentials to firing probabilities is non-linear, which is one of the main differences between SGP CaKe and the Hawkes process. Similarly, the rightmost figures show how the firing rates may be reconstructed using the variational method.

Figure 2 shows the variational approximation of the membrane potential for the first network, this time for different coupling strengths *w*. As the figure shows, the membrane potentials recovered well for the neurons that received no input (i.e. the top row of the figure). For the neurons that did receive input the membrane potential approximation deteriorates in its estimation of the magnitude when the coupling strength is increased. This is due to the violation of the assumption that neurons are only weakly coupled and have their activity predominantly driven by internal dynamics. Despite this, the correlation between the true and the estimated membrane potential remains high and, as we will show below, sufficient to recover the causal structure.

**Figure 2:**
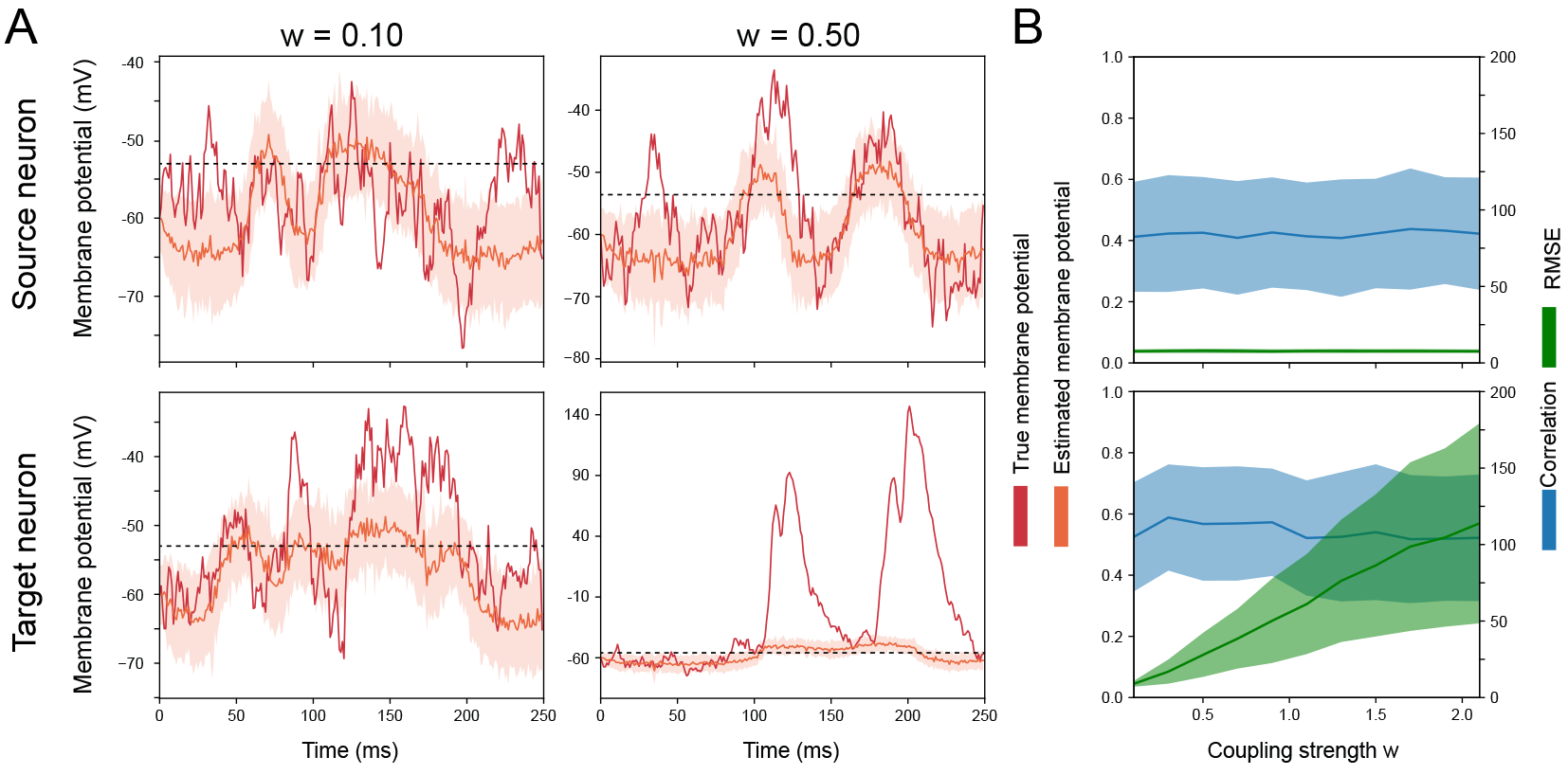
**A.** Variational inference of the membrane potentials for the source and target neurons of network 1 (see Fig. 4A, for *w* ∈ {0.1,0.5}). The dashed line indicates the membrane threshold. **B.** The correlation and root-mean-squared error between the true and estimated membrane potentials. Shaded intervals indicate one standard deviation.

### 5.1 Recovery of effective connectivity

As an example, Fig. 3 shows the recovered causal response functions for the two-neuron network with a single inhibitory connection. Both variants of SGP CaKe successfully distinguish present and absent coupling and correctly identify that the present connection is inhibitory. In addition, we show the cross-correlation estimation of this connection. While this more traditional approach also identifies the inhibitory coupling, it fails to classify the other connection as absent, as can be seen from the estimated effect sizes (see Appendix C) for the two connections.

**Figure 3:**
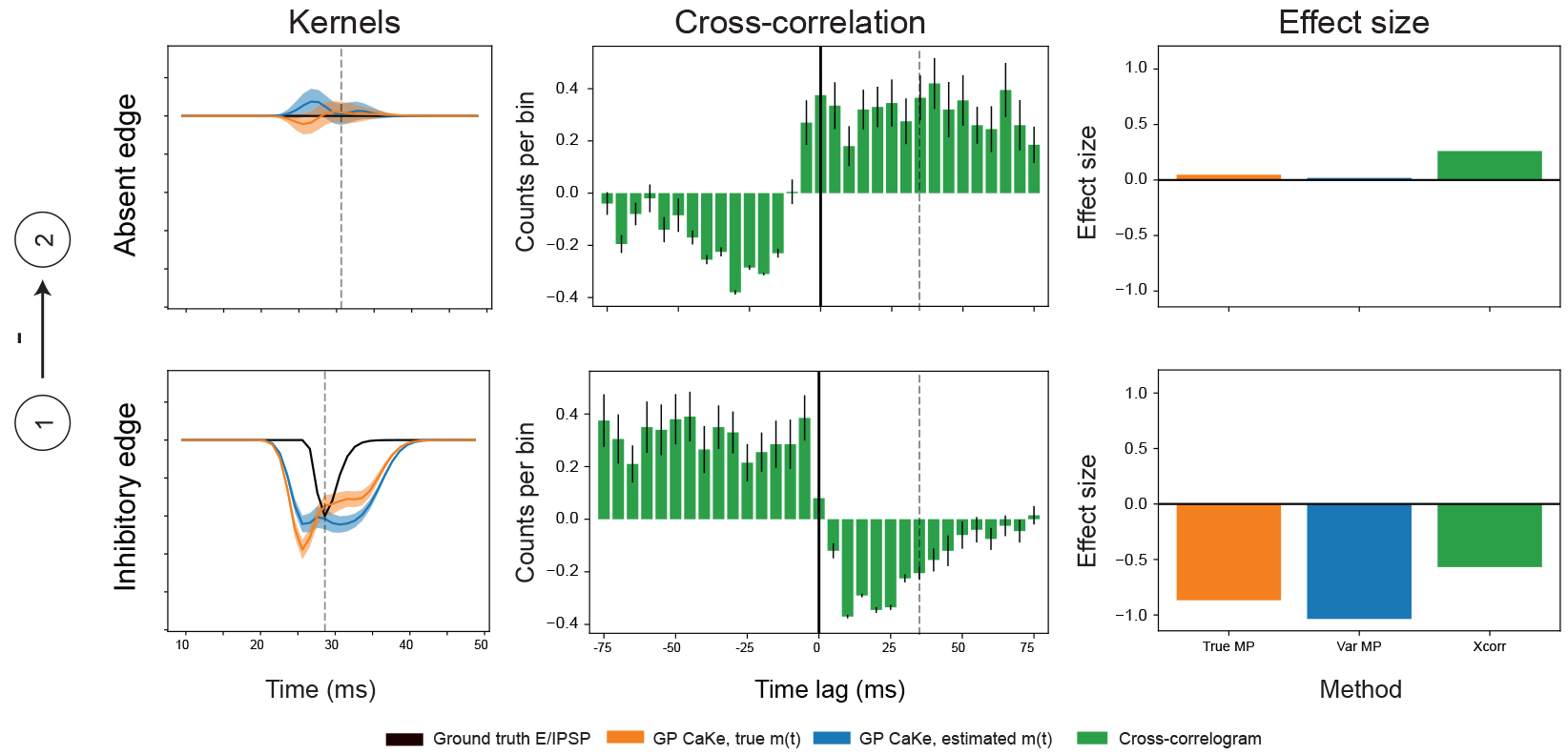
Examples of reconstructed causal kernels for both variants of SGP CaKe together with the corresponding cross-correlograms and effect sizes for each of the three methods, per connection. Shaded intervals and error bars indicate 95% confidence intervals.

To further quantify these results we use the estimated causal response functions to recover the coupling structures from Fig. 4A. The presence of a connection is estimated via a *z*-test at the peak of the true causal response function while its directionality is given by the sign of the corresponding z-score (more details are provided in Appendix C). The performance is scored using the root-mean-squared-error between the true adjacency matrix describing the coupling structure and the estimated structure. The results of this analysis are shown in Fig. 4B. From these results it is apparent that both SGP CaKe variants consistently provide the best estimates of the causal coupling structure. In many cases the recovery is (near) perfect.

**Figure 4:**
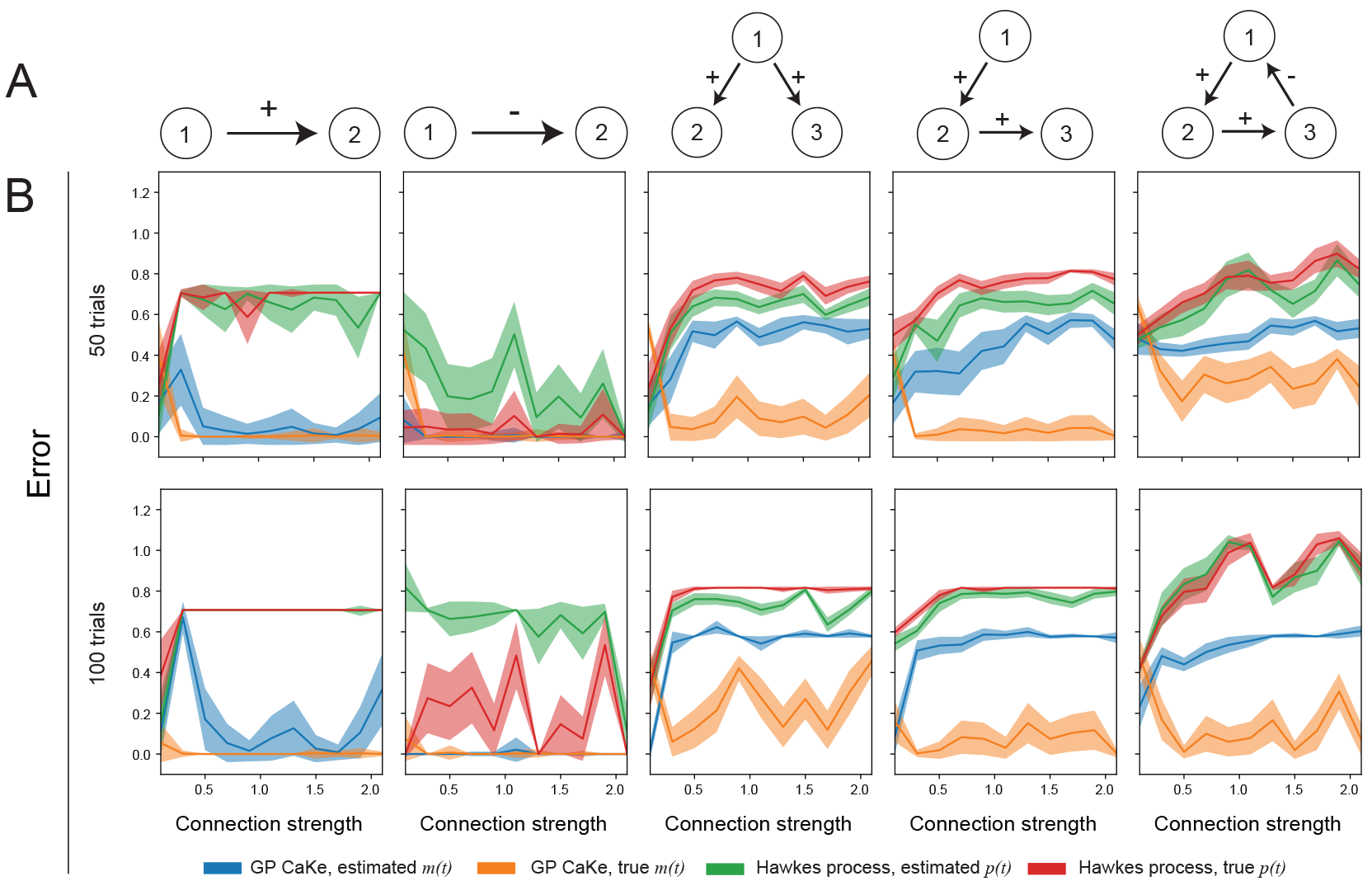
**A.** The considered network structures. **B.** Root-mean-squared-error between the actual connectivity matrix and the different recovery approaches shown for different numbers of bootstrapped subsamples. Interval widths indicate one standard deviation over 1 000 runs of the indicated number of bootstrapped samples.

When the coupling strength *w* is increased the assumption of weak coupling is again violated. We see that this is particularly detrimental for the networks with common causes and transitive effects. However, when the true membrane potential is observed, SGP CaKe still estimates the coupling structure nearly perfectly. Also, even for these more complex cases, the variational SGP CaKe approach outperforms both variants of the Hawkes process, even the idealized case where the true firing rates are known. Interestingly, for some networks the Hawkes process in fact performs better with the estimated firing rates than with the true ones. Presumably this is due to the smoothing induced by the variational approach, which causes the estimated firing rates to be more similar to membrane potentials.

## 6 Analysis of real spike trains: connectivity in rat entorhinal cortex

To illustrate a more realistic application of our proposed methods we applied both the SGP CaKe and the Hawkes process to multi-unit recordings of rat entorhinal cortex [26, 27]. Details of the data acquisition and preprocessing can be found in Appendix D. For these data sets only the spikes were observed so we estimated the membrane potentials and firing rates for SGP CaKe and the Hawkes process respectively using the semi-analytic variational approach. As the ground truth is obviously unavailable, we estimated coupling in two different conditions (condition one consists of the rat moving freely in an open square; condition two consists of the rat navigating through a linear maze) and looked at the between-session reproducibility for validation of the procedures.

The estimated causal response functions between three electrodes are shown in Fig. 5. Overall, SGP CaKe and the Hawkes process resulted in similar causal response functions, although slight differences may be observed in the estimated coupling structure. Clearly there is strong correspondence in the causal response functions within conditions, while at the same time the response functions between the conditions are fairly different, showing the sensitivity of the methods. The reproducibility is further quantified in the correlations between the causal response functions for each pair of conditions (see inset in Fig. 5).

**Figure 5:**
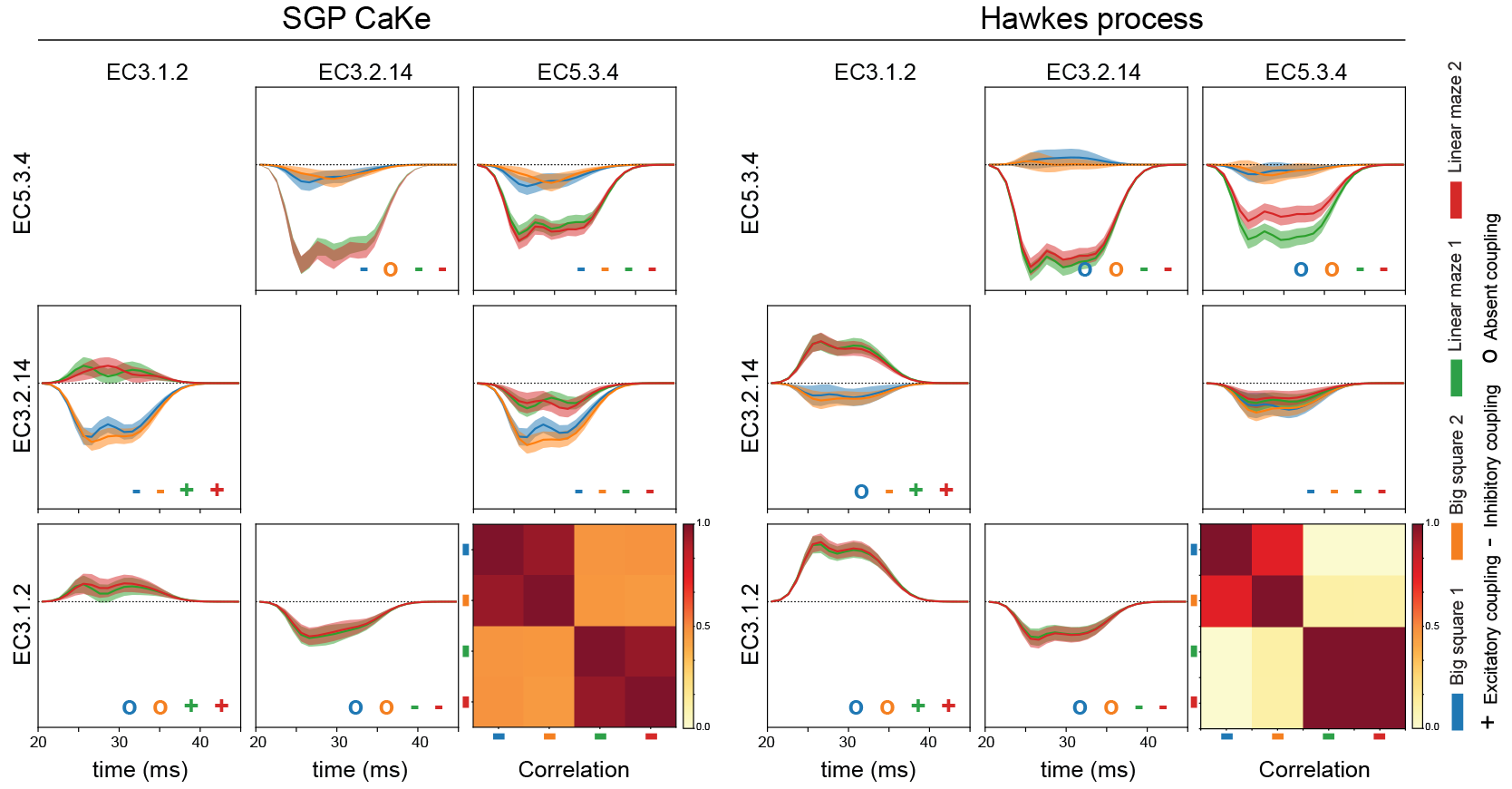
Estimated causal connectivity between three neuronal clusters in entorhinal cortex for four conditions (see Appendix D. The insets show the (average) correlation between the causal response functions for the four different conditions

## 7 Conclusion

We introduced two new nonparametric Bayesian models for spike-membrane and spike-spike connectivity analysis. We obtain an approximate semi-analytic posterior for the spike-spike problem by minimizing a new likelihood-free variational loss. This semi-analytic method has wide applicability outside our current model since it can be used every time a latent GP regression is coupled to a non-linear emission model. For example, our semi-analytic variational method can be directly used in a calcium imaging setting where the spikes are observed through a non-linear calcium response [28].

